# Neuronal LXR Regulates Neuregulin-1 Expression and Sciatic Nerve-Associated Cell signaling

**DOI:** 10.1101/2020.01.14.907048

**Authors:** Chaitanya K. Gavini, Raiza Bonomo, Virginie Mansuy-Aubert

**Author notes:** Corresponding author: Virginie Mansuy-Aubert; Cell and Molecular Physiology, Stritch School of Medicine, Loyola University Chicago, Maywood, Illinois, USA 60153.

## Abstract

Neuropathic pain caused by peripheral nerve injury significantly affects sensory perception and quality of life. Accumulating evidence strongly link cholesterol and inflammation with development and progression of Obesity and Diabetes associated-neuropathies. However, the exact mechanisms of how lipid metabolism in peripheral nervous system (PNS) contributes to the pathogenesis of neuropathy remains poorly understood. Dysregulation of LXR pathways have been identified in many transcriptomic analyses in neuropathy models. LXR α/β expressed in sensory neurons are necessary for proper peripheral nerve function. Deletion of LXR α/β from sensory neurons lead to pain-like behaviors. In this study, we identified that LXR α/β expressed in sensory neurons regulates neuronal neuregulin-1 (Nrg1). Using in vivo cell-specific approaches, we observed that loss of LXR from sensory neurons altered genes regulating lipid metabolism in non-neuronal cells potentially representing Schwann cells (SC). Our data suggest that neuronal LXR may regulates SC function via a Nrg1-dependent mechanism. The decrease in Nrg1 expression in DRG neurons of WD-fed mice may suggest an altered Nrg1-dependent neuron-SC communication in Obesity. The communication between neurons and non-neuronal cell such as SC could be a new biological pathway to study and to treat Obesity-associated neuropathy and PNS dysfunction.

## Introduction

Peripheral neuropathy arising from metabolic disorders such as Obesity and pre-diabetes leads to loss of sensory perception, pain, and reduced quality of life^1-3^. Currently, there are no disease modifying drugs available, as the neurobiology underlying the neuropathic pain is still unclear. Many previous studies have focused on glucose as a major culprit in the development of Obesity-associated neuropathies ^4-6^, but recent body of evidence indicated that altered plasma lipid signaling in the peripheral nervous system (PNS) ^7-10^ is involved. Indeed, evidence on obese and diabetic patients, and murine model of Type II diabetes strongly suggest that circulating cholesterol and inflammation are linked to development and progression of neuropathy^11-13^. However, the mechanisms underlying how lipids affect cells of the PNS are still unclear. Notably, sensory neurons located in the dorsal root ganglia (DRG) and associated Schwann cells (SC), satellite cells or immune cells in the nerves are not protected from the blood-brain or blood-nerve barrier^14^ suggesting that, in contrast to the central nervous system (CNS), they could sense and/or be affected by circulating lipids. Previously, we have found that peripheral neurons express liver X receptors (LXR α/β) among other lipid sensors^15-17^. LXR α/β are ligand-activated nuclear receptors that bind metabolites of cholesterol^18,19^. Many previous findings strongly suggested that, LXRs play an important role in nervous systems^16,17^ but, while the function of LXR α/β in regulating cholesterol efflux (in liver, intestine, adipose tissue or macrophages) and triglycerides synthesis (in liver) is well characterized, its specific cellular role in the PNS remains understudied.

In the peripheral system, neurons are associated with SC and they have a high interdependent relationship i.e., damage to one cell type leads to pathophysiological changes in the other. SC are under the control of axons and their nuclei located in DRG^20-22^. The SC population is heterogeneous since some SC produce myelin and others do not, however, in all cases reported, SC are crucial for normal PNS function and repair ^23-25^. These specific roles are understudied *in vivo* because markers defining specific SC population are unknown. Membrane-bound neuregulin-1 type III (Nrg1-type III) is expressed in neurons and regulates neuron-associated SC functions including the formation or no of myelin sheath^26^. Expression of Nrg1-type III, independent of axon diameter, provides the signal that determines whether axons become unsheathed, myelinated or repaired after nerve injury^27-29^. Neuronal Nrg1 interaction with SC epidermal growth factor receptors (ErbB) is necessary to maintain normal peripheral nerve function^30^. Importantly, disrupting interaction of Nrg1 and ErbB at the SC–axon interface leads to abnormal and aberrant myelination of large fibers, and perturbation in the Remak bundles structure containing small non-myelinated axons, resulting in abnormal nerve conduction and altered nerve structure^30,31^. While neuronal Nrg1/ErbB expression is crucial to regulate SC function, its gene regulation is unclear either in normal or pathological states. Our study using mice models of diet-induced Obesity suggest that neuron-non-neuronal cell (potentially SC) communication is altered in PNS of diet-induced Obesity. *In vivo* cell-specific approaches demonstrate that LXRs expressed in nociceptors regulate *Nrg1* expression, and modulate lipid homeostasis in cells of the nerve likely in a paracrine manner, unmasking a unique pathway that could be targeted to improve neuropathy associated with Obesity.

## Results

### Decreased ErbB expression in the nerves of western diet-fed mice

Others and we identified that WD-feeding led to peripheral neuropathy affecting the sensory neurons and their associated SC^16,32^. Previous reports demonstrated that modification in SC metabolism in the sciatic nerve were strongly linked to neuropathy phenotype in obesity^3^. To identify changes in transcriptome in SC after WD, we processed sciatic nerve of normal chow (NC) or WD-fed wild type mice for RNA sequencing (RNA-seq) after 14 weeks of diet and analyzed for differentially regulated genes and pathways. Sciatic nerves from NC- and WD-fed mice were dissected, total RNA was purified and samples that passed quality test were subjected to RNA sequencing (Figure1A). Sensory neurons are associated with both non-myelinating and non-myelinating SC, to our knowledge there are no markers that define the entire population (non-myelinating and myelinating). The sciatic nerve is comprised of various cells types such as immune cells, satellites cells, but SC are known to represent the major cell type in the sciatic nerve and partners with all axons providing metabolic support ^33,34^ suggesting that most of gene identified in RNAseq data are SC genes. In the future, specific markers would need to be identified to confirm this statement.

**Figure1:**
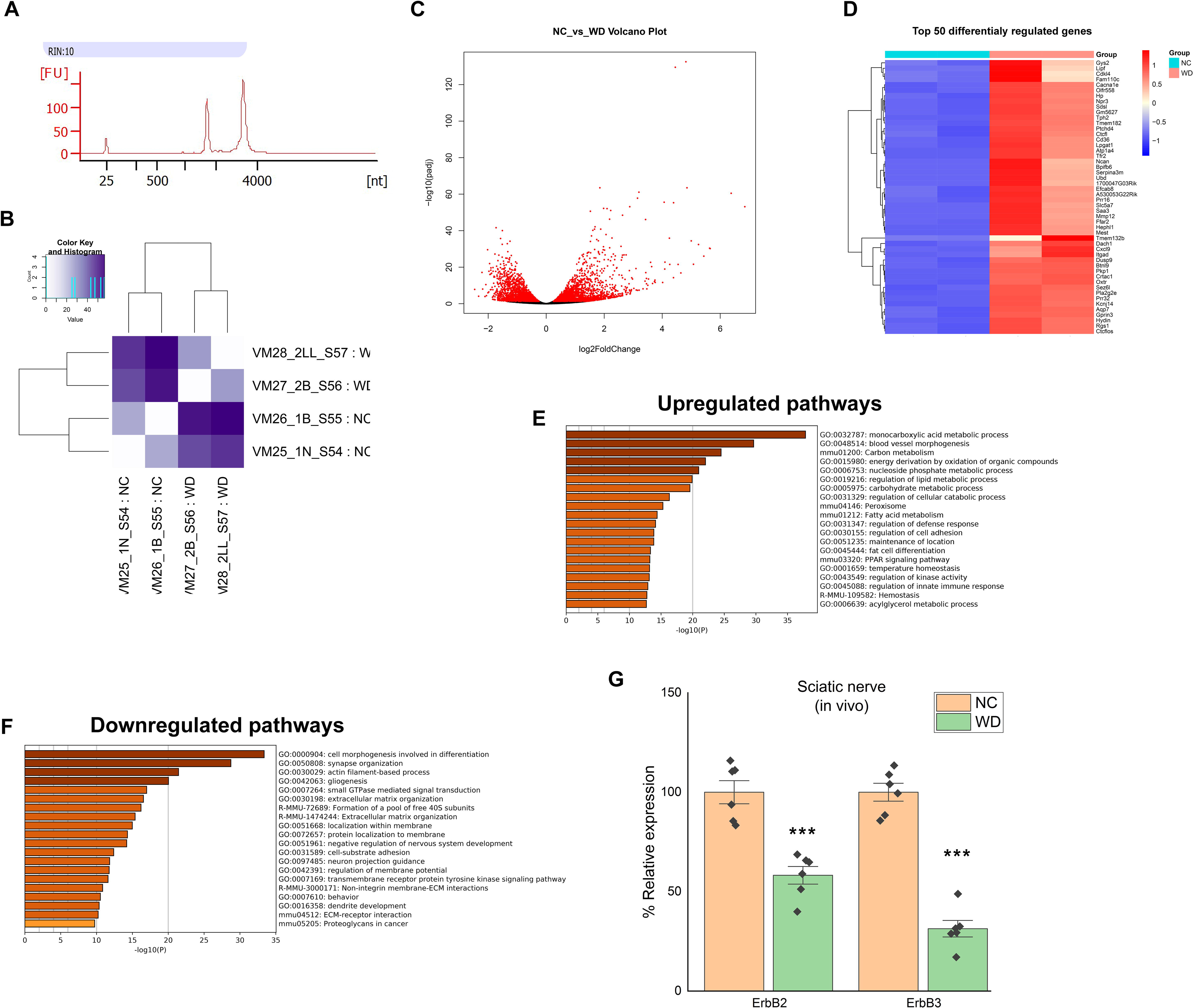
WD-fed mice have downregulation of axon guidance and Schwann cell homeostasis pathways in sciatic nerve. A) RNA quality assessed by Agilent Bioanalyzer using Total RNA Pico Chip. B) Heat map showing the PCA distances between each biological replicate. C) Volcano plot revealing upregulated and downregulated genes in the transcriptome (n=2 biological replicates). D) Heat map of top 50 differentially regulated genes in sciatic nerve of NC or WD-fed mice (n=2 biological replicates). E) Pathway analysis of upregulated genes from RNA-seq in sciatic nerve of NC or WD-fed mice (n=2 biological replicates). F) Pathway analysis of downregulated genes from RNA-seq in sciatic nerve of NC or WD-fed mice (n=2 biological replicates). G) qPCR verification of mRNA level of erbb2 and 3 in sciatic nerve of NC or WD-fed mice with NC-fed group treated as 100% (n=6/group). All data are Mean±S.E.M. ***p<0.0005

RNA-seq analysis identified 4,166 differentially regulated genes between NC and WD-fed mice sciatic nerves (Figure1 B, C, and D). 2153 genes were upregulated and 1970 genes were downregulated in the sciatic nerve of WD-fed mice (Figure1C). To identify pathways that were significantly enriched in upregulated and downregulated genes after WD, we performed Metascape analysis. Metascape pathway analysis revealed significant enrichment of genes involved in lipid and carbohydrate metabolism-related pathways including “lipid membrane homeostasis” (e.g. Cd36, Abcg1, Abca1, Pparγ) in upregulated dataset (Figure1E). Many of the genes dysregulated were consistent with previous studies in others models of diabetic neuropathy (e.g. Abca1, Abcg1, Wnt, Srebpf1) ^35-38^. In addition, we observed enrichment of gene involved in “PNS function” “cell survival”, “migration”, “axon guidance”, and “receptor interaction and signal transduction-related” pathways (ErbB2 and 3, Slit2, Sox10, Sirt2, Fgfr, Piezo 2) (Figure1F). Interestingly, ErbB genes are implicated in SC function and are known to be necessary to maintain normal peripheral nerve function^30^ via a neuronal-SC communication. For the first time, we identify this pathway dysregulated in the nerve of 14 weeks WD-fed mice. To validate decrease in ErbB2, and ErbB3, qPCR analyses were performed in independent experiments using separate cohorts of mice. We confirmed significant decrease in mRNA levels of genes involved in ErbB signaling pathway (ErbB2 and 3) in sciatic nerve of WD-fed mice compared to NC-fed (Figure1G). Our data suggest that the communication between neuron and SC may be altered in the PNS of WD-fed mice compared to NC mice.

### NRG1 expression is decreased in the DRG of western diet-fed mice

As mentioned, DRG axons are in continuous contact with non-myelinating and myelinating SC. Axonal cues, in particular NRG1, is known to be a driving force for regulating SC proliferation and migration^26^. Since neuronal Nrg1-type III regulates neuron-SC communication involving SC ErbB pathways^26^, we assessed the level of Nrg1-type III in the DRG (where neuronal cell bodies are located) of NC versus WD-fed mice. We observed that DRG of WD-fed mice have a significantly lower mRNA level of Nrg1-type III compared to NC-fed mice, (Figure2A). To evaluate whether the decrease of Nrg1-type III expression in WD-fed mice may be due to an effect of saturated fat on the neurons, we used *ex vivo* and *in vitro* approaches. First, organotypic cultures of DRG were stimulated with the saturated fatty acid, palmitic acid (400µM). Compared to vehicle (BSA)-treated DRG, palmitate-treated DRG had lower mRNA level of Nrg1-type III (Figure2B). As DRG contains several cell types, second, we dissociated DRG to study Nrg1 in primary neuron cultures. Compared to vehicle treated, neurons exposed to palmitic acid had also lower mRNA level of Nrg1-type III (Figure2C) suggesting that saturated fat can alter the expression of Nrg1-type III and potentially neurons/SC communication.

**Figure2:**
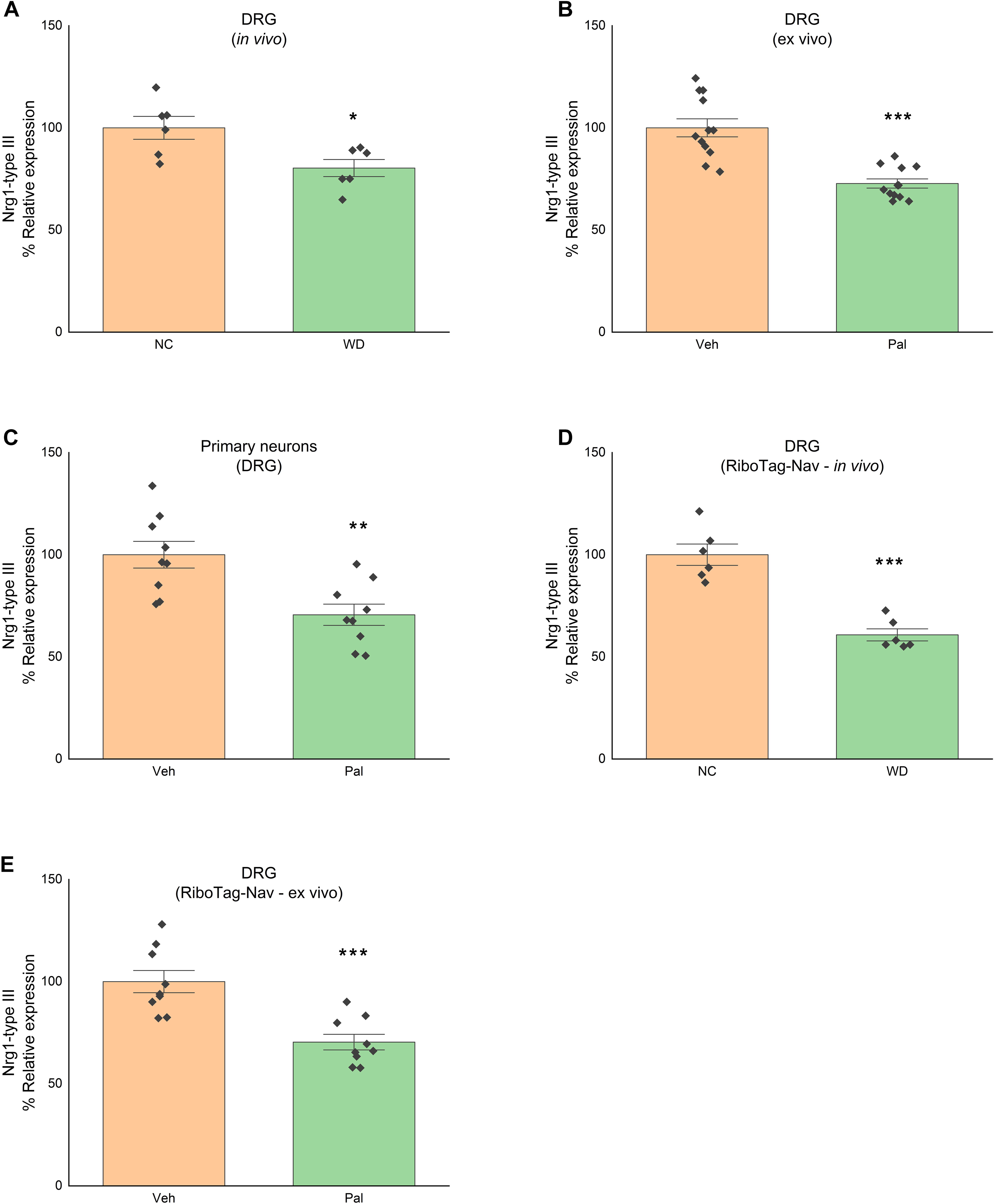
Nrg1-type III level is decreased in DRG sensory neurons of WD-fed mice. A) Nrg1-type III mRNA level in DRG of mice fed either NC or WD with NC group treated as 100% (n=6/group). B) Nrg1-type III mRNA level in organotypic culture of DRG stimulated with or without palmitate (vehicle group treated as 100%) (n=4 experiments in triplicate). C) Nrg1-type III mRNA level in primary neuronal culture of DRG stimulated with or without palmitate (vehicle group treated as 100%) (n=3 experiments in triplicate). D) Nrg1-type III mRNA level in DRG sensory neurons of RiboTag-Nav mice fed either NC or WD with NC group treated as 100% (n=6/group). E) Nrg1-type III mRNA level in organotypic culture of DRG sensory neurons (from RiboTag-Nav mice) stimulated with or without palmitate (vehicle group treated as 100%) (n=3 experiments in triplicate). All data are Mean±S.E.M. *p<0.05; **p<0.005; ***p<0.0005

Previous studies including ours have shown that Nav1.8 expressing neurons in the DRG play a critical role in the pathogenesis of WD-associated neuropathy^16,32^. To specifically test, the effect of WD or of saturated fatty acids on Nrg1 expressed in sensory neurons of the DRG, we directly purified mRNA in translation (associated with ribosomes) from DRG sensory neurons. To this end, we generated mice expressing a HA-tagged ribosomal protein (RPL22-HA) in the sensory neurons (RiboTag+/+:Nav1.8Cre+/-; RiboTag-Nav) by crossing RiboTag mice with hemizygous Nav1.8-Cre mice^16^. Sensory neuron specific mRNAs and in translation were isolated from DRG of RiboTag-Nav as described before^16^. Compared to NC-fed mice, sensory neurons of WD-fed (RiboTag-Nav) mice have less Nrg1-type III mRNA in translation (Figure2D). Similar results were obtained from e*x-vivo* DRG organotypic cultures isolated from RiboTag-Nav and treated with palmitate (Figure2E). These results along with previous studies^16^ suggest saturated lipids overload during WD, impairs Nrg1 expression and likely the communication between sensory neurons and others cell types, likely SC.

### LXR are transcriptionally active in DRG and regulates Nrg1 expression

Next, we sought to determine the cause of the decrease in Nrg1-type III in sensory neurons of WD-fed mice, and following saturated fatty acid stimulation. Interestingly, Nrg1-activated ErbB pathway had been shown to regulate cholesterol biosynthetic pathway in a paracrine manner ^39^. LXRs (LXRα and LXRβ), are lipid nuclear receptors, and play a crucial role in regulation of cholesterol and fatty acid homeostasis ^40^. Our previous study using high-throughput real-time PCR screen showed that LXRs are expressed in DRG and LXR agonist treatment prevents progression of obesity-induced allodynia^16^. First, using *in-situ* hybridization, we confirmed the presence of LXRs transcripts in the DRG (Figure3A). To directly test the transcriptional activity of LXR in DRG neurons, we transduced primary DRG neurons with a pGreenFire1-LXRE Lentivector reporter that co-expresses a destabilized copepod GFP and luciferase from the LXR response elements and neighboring regions in the LXRα promoter (Figure3B). Activity of LXR was evaluated by measuring the change in GFP expression between groups as well as using luciferase assay. Stimulation of transduced DRG neurons with GW3965, a potent LXR agonist, significantly increased the overall GFP fluorescence measured by flow cytometry (Figure3C) along with increased luminescence during luciferase assay (Figure3D). We also evaluated control canonical targets of LXR α/β implicated in cholesterol efflux (Abca1), cholesterol synthesis (Srebf1/Srebp-1c), and confirmed that their expressions were increasing following LXR activation (Figure3E). These data demonstrate that LXRs are transcriptionally active in DRG neurons. To test the activity of LXRs in the presence of saturated fatty acid, transduced neurons with the lentivector reporter were treated with palmitic acid followed by GW3965 stimulation. Palmitate treatment significantly lowered the increase in luminescence induced by GW3965 (Figure3D). These findings confirm that LXR activity and its canonical gene expressions are altered in Obesity in neurons as in many other tissues^16,40,41^.

**Figure3:**
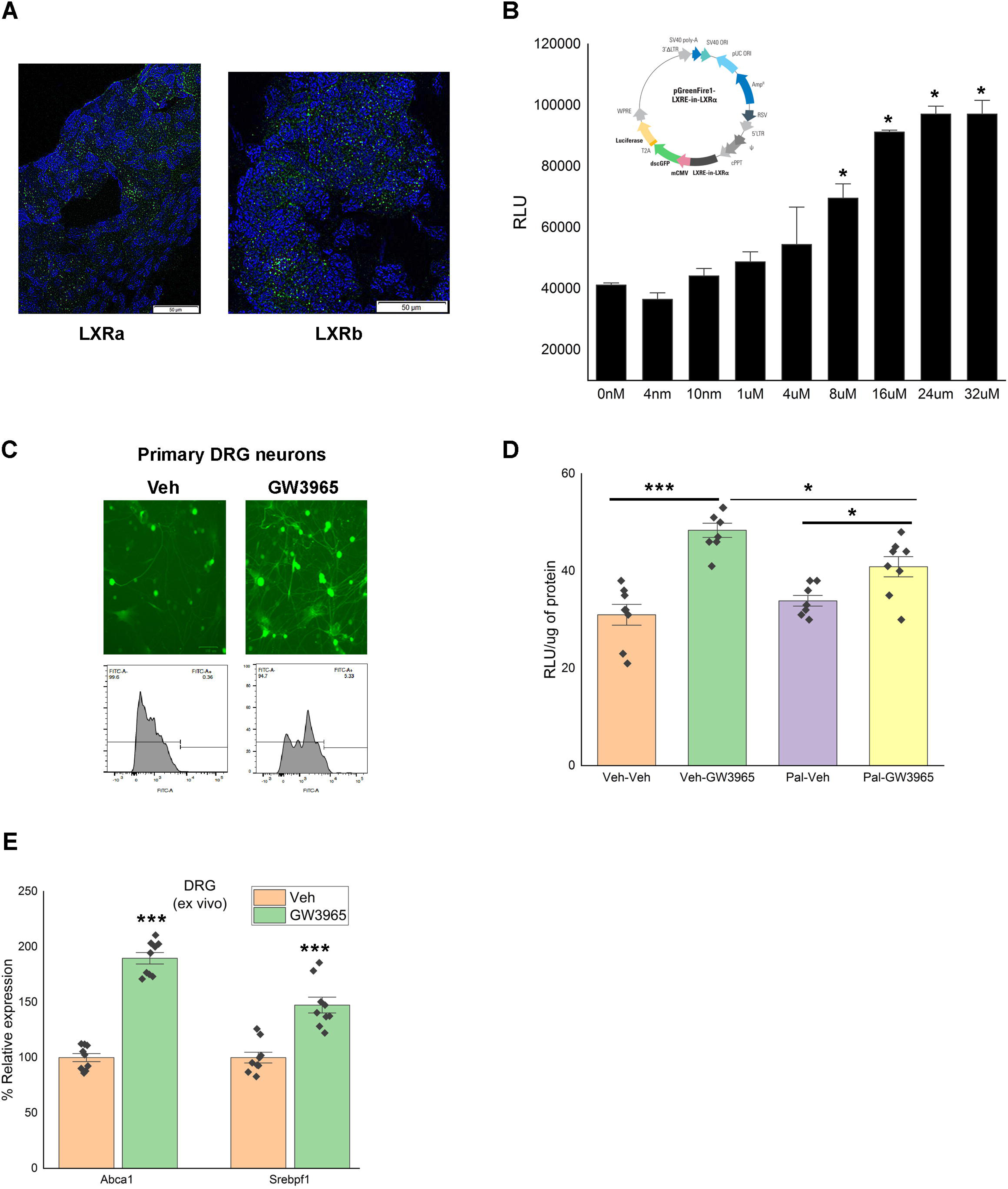
LXR are transcriptionally active in DRG. A) In situ hybridization assay confirming presence of LXRa and LXRb mRNA in mouse DRG (scale 50µm). B) Luminescence dose-response curve for LXR agonist, GW3965, in DRG primary neuronal cultures transduced with pGreenFire1-LXRE in LXRα Lentivector (n=3 individual experiments) (insert - pGreenFire1-LXRE in LXRα Lentivector). C) GFP expression of GW3965 treated DRG primary neurons transduced with pGreenFire1-LXRE in LXRα Lentivector using fluorescence imaging and flow cytometry (n=3 individual experiments) (scale 100µm). D) GW3965 mediated-luciferase activity in DRG primary neuronal cultures transduced with pGreenFire1-LXRE in LXRα Lentivector treated with or without palmitate (n=3 experiments in triplicate). E) Activation of LXR by GW3965 increases gene expression of LXR canonical pathway in the DRG (vehicle group treated as 100%) (n=3 experiments in triplicate). All data are Mean±S.E.M. *p<0.05; ***p<0.0005

Next, we hypothesized that LXR could regulate Nrg1 gene expression. To evaluate the effect of LXR activation on the Nrg1-type III expression in normal and in an obesogenic environment, we treated mice fed either NC or WD for 8 weeks with LXR agonist, GW3965 (25mg/Kg of body weight) twice a week for 3 weeks. Compared to NC-fed mice, DRG of WD-fed mice had lower level of Nrg1-type III mRNA. GW3965 administration increased Nrg1-type III mRNA level in the DRG of WD-fed mice (Figure4A). To directly assess whether GW3965 can affect the DRG neurons, we stimulated DRG organotypic cultures and primary DRG neuronal cultures with GW3965 and palmitate (Figure4B, C). Compared to vehicle, GW3965 treatment significantly increased mRNA expression of DRG Nrg1-type III that was blunted in presence of palmitate (Figure4B, C).

To test whether GW3965 regulates Nrg1-type III expression in sensory neurons, NC or WD-fed RiboTag-Nav mice or palmitate-stimulated DRG organotypic cultures from RiboTag-Nav mice were treated with GW3965 as describe above. Compared to NC-fed RiboTag-Nav, sensory neurons of WD-fed RiboTag-Nav mice have lower mRNA level of Nrg1-type III (Figure4D, E) and GW3965 treatment attenuated this decrease (Figure4D, E). Overall data suggest that LXR α/β regulate Nrg1 expression in the sensory neurons of the DRG. Our findings also suggest that the decrease in Nrg1 mRNA level observed *in vivo* in the nociceptors of obese mice i) may be the consequence of a decrease in LXR activity(induced by diet and/or saturated fatty acids) and ii) could be rescued by GW3965.

**Figure4:**
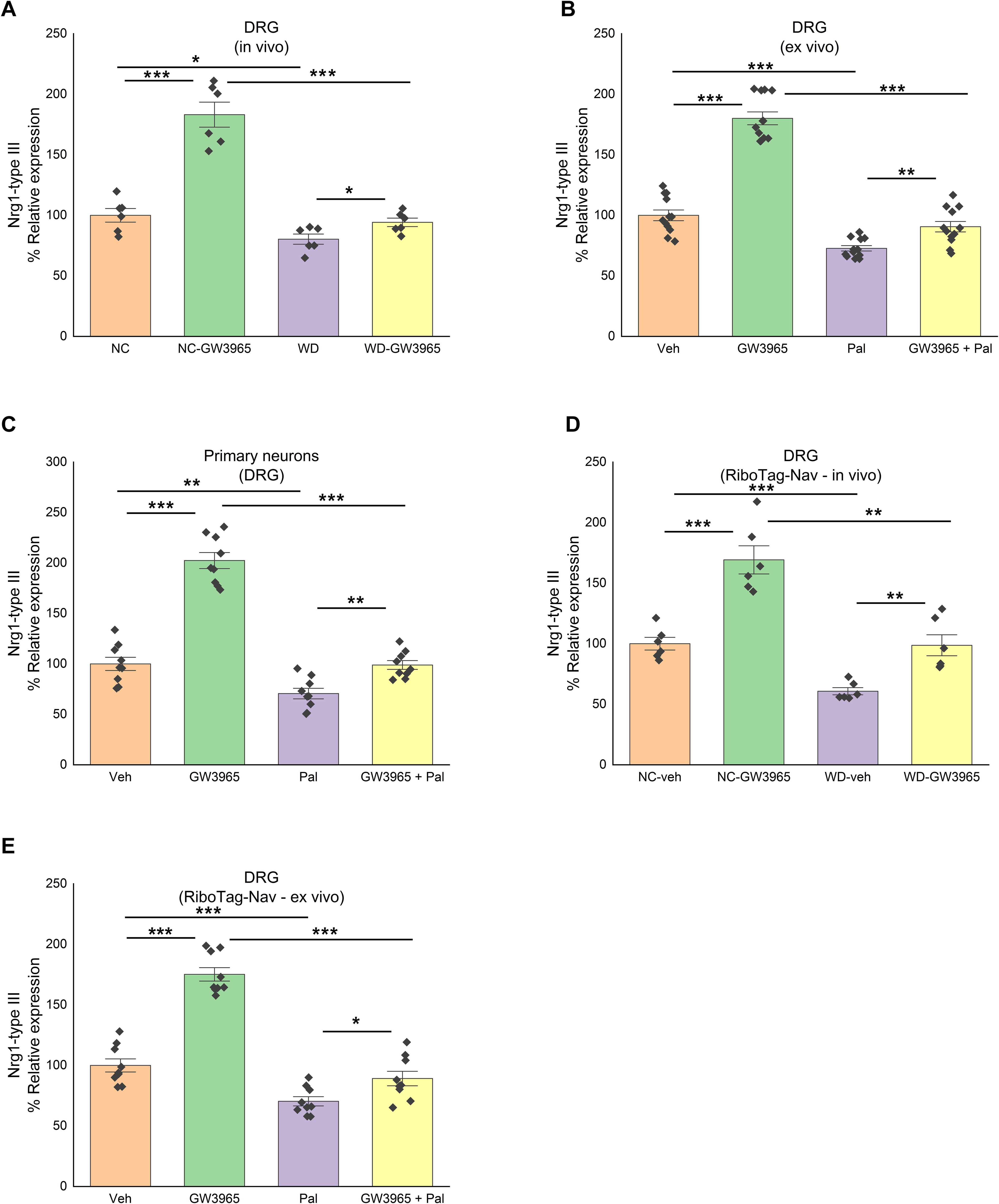
LXR activation increases Nrg1-type III expression in sensory neurons of the DRG. A) Nrg1-type III mRNA level in DRG of mice fed either NC or WD and treated with or without GW3965 with NC group treated as 100% (n=6/group). B) Nrg1-type III mRNA level in organotypic culture of DRG stimulated with or without palmitate and treated with or without GW3965 (vehicle group treated as 100%) (n=4 experiments in triplicate). C) Nrg1-type III mRNA level in primary neuronal culture of DRG stimulated with or without palmitate and treated with or without GW3965 (vehicle group treated as 100%) (n=3 experiments in triplicate). D) Nrg1-type III mRNA level in DRG sensory neurons of RiboTag-Nav mice fed either NC or WD and treated with or without GW3965 with NC-veh group treated as 100% (n=6/group). E) Nrg1-type III mRNA level in organotypic culture of DRG sensory neurons (from RiboTag-Nav mice) stimulated with or without palmitate and treated with or without GW3965 (vehicle group treated as 100%) (n=3 experiments in triplicate). All data are Mean±S.E.M. *p<0.05; **p<0.005; ***p<0.0005

### Loss of LXRs in sensory neurons decrease neuronal Nrg1 expression and change lipid gene expressions in the nerve cells potentially SC

Our current data suggests that loss of LXRs activity in the sensory neurons of WD-fed mice could decrease Nrg1 expression impairing neuron-Schwann cell communication. This would result in impairment of ErbB downstream pathways in SC. To test this *in vivo*, we generated sensory neuron specific deletion of LXRs mouse model (LXRαfl/flβfl/fl:Nav1.8Cre+/-; LXRabnav) by crossing LXRαfl/flβfl/fl (LXRab) mice with Nav1.8Cre+/- mice as previously reported (Figure5A)^16,17^. LXRab and LXRabnav mice were fed either NC or WD and their DRG were processed to assess the mRNA levels of LXR canonical pathway and Nrg1. As shown in Figure5B, Abca1 was decreased more than two fold in both NC and WD-fed LXRabnav mice. Srebf1/Srebp-1c were decreased in DRG of WD-fed mice but in contrast to abca1, no difference was observed between LXRab and LXRabnav suggesting that, in sensory neurons, LXR does not drive Srebf1/Srebp-1c. Interestingly, Nrg1-type III was significantly reduced when LXRs are absent from sensory neurons demonstrating that endogenous LXRs regulate Nrg1-type III expression in sensory neurons *in vivo* and that WD feeding may alter this regulation (Figure5B).

**Figure5:**
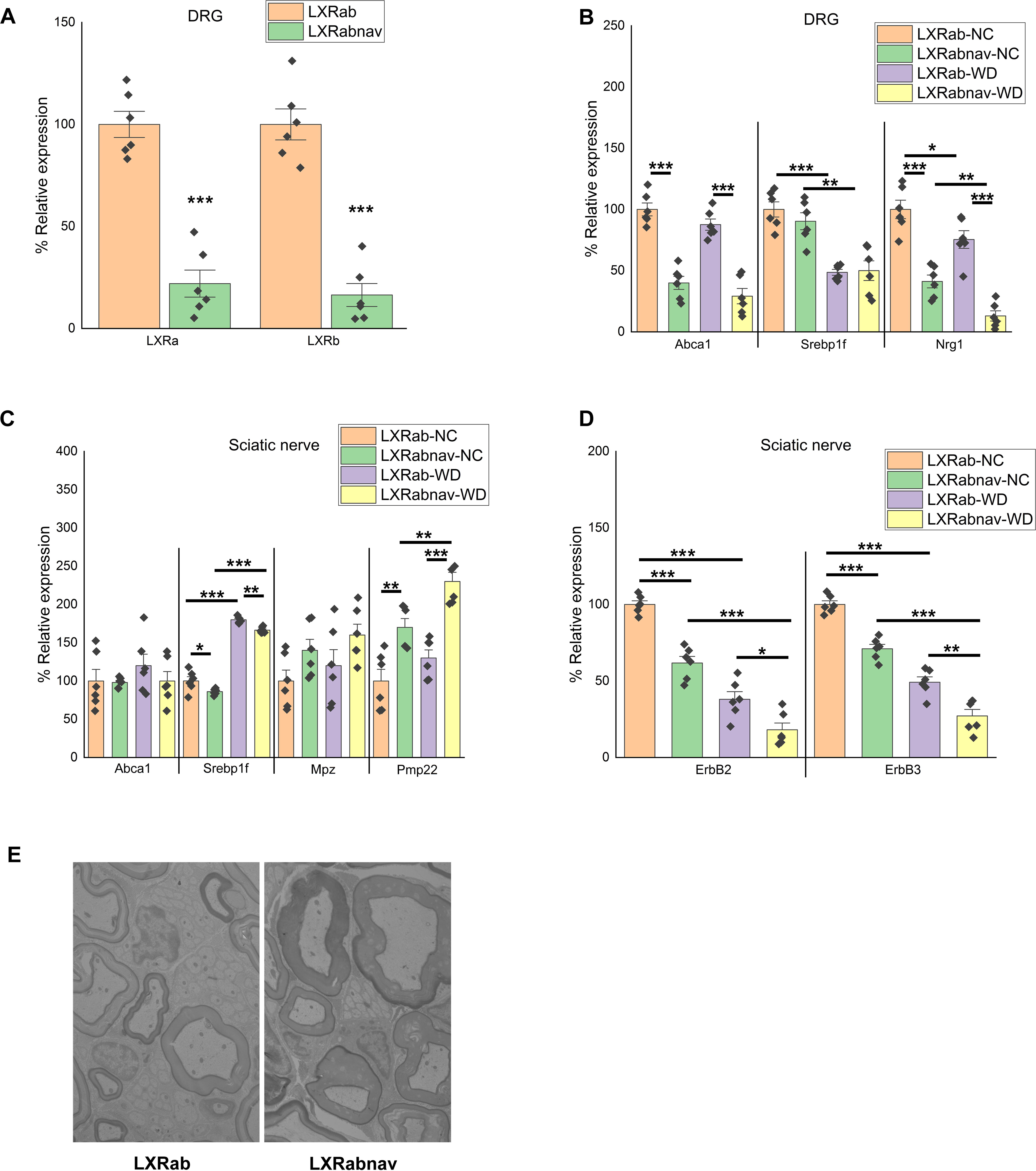
Loss of LXR α/β in sensory neurons decrease neuronal nrg1 expression. A) LXRα and LXRβ mRNA levels in DRG of LXRα/β cell specific knockout using Nav1.8-Cre with control group set at 100% (n=6/group). B) mRNA levels of LXR canonical targets in the DRG of LXRab and LXRabnav mice fed either NC or WD (with LXRab-NC group set to 100%) (n=6/group). C) mRNA levels of LXR canonical targets in the sciatic nerve of LXRab and LXRabnav mice fed either NC or WD (with LXRab-NC group set to 100%) (n=6/group). D) B) mRNA levels of erbb2 and 3 in the sciatic nerve of LXRab and LXRabnav mice fed either NC or WD (with LXRab-NC group set to 100%) (n=6/group). E) Electron microscopic images of sciatic nerve from LXRab and LXRabnav mice. All data are Mean±S.E.M. *p<0.05; **p<0.005; ***p<0.0005

As mentioned above, DRG contains cell bodies of axons that compose the sciatic nerve and are associated with myelinating or non-myelinating SC. As Axonal Nrg1 is known to determine SC function^26,27^, we isolated sciatic nerves from LXRab and LXRabnav mice to assess gene expression involved in lipid metabolism that were previously shown to be downstream of Nrg1/ErbB^25-27,30^: Srebf1/Srebp-1c, myelin protein zero (MPZ), and peripheral myelin protein (PMP22) were evaluated. We rigorously dissected the nerve to harvest the same piece of tissue in each experiment. In pilot data, the number of cells isolated were counted, the total number of total cells were similar between all experiments (not shown). Interestingly, Srebf1/Srebp-1c expression increased in WD-fed mice sciatic nerve compared to NC-fed mice (Figure5C). We did not observe any differences between genotypes suggesting that Srebp1f change following WD in nerve-associated cells is independent of neuronal LXR (Figure5C). We observed an increase of Mpz and Pmp22 mRNA in sciatic nerve of LXRabnav mice compared to control mice. Compared to LXRab, we also observed a decrease in ErbB2 and ErbB3 mRNA in the sciatic nerve of LXRabnav mice (Figure5D). This decrease was exacerbated when fed WD (Figure5D). These data suggest loss of LXR in sensory neurons may affect signaling in nerve-associated cells and may change myelin structure. We evaluated the sciatic nerve of LXRab and LXRabNav mice using electron microscopy, but we did not see any significant changes in SC number or structure in either Remak bundles (non-myelinated SC) or myelinating SC associated with the nerve (Figure5E). It is also possible that the change in gene expression is seen at an earlier time point than that of the change in structure or that the EM fails to capture these changes. Cell-specific models (e.g -cre) allowing to specifically purify SC non-myelinating and myelinating would be necessary to quantify *in vivo* the numbers of all SC and also to directly purify mRNA. Unfortunately, these models do not exist, the current knowledge on SC rely on *in vitro* experiment using SC cultures. These changes in neuron-nerve associated cell signaling may be involved - at least in part-in the mechanical and thermal hypersensitivity observed in obese mouse model lacking LXR in sensory neurons (as published^16,32^).

## Discussion

Human studies showed correlation between circulating cholesterol and lipids in development and progression of neuropathy^11-13^ albeit the mechanisms remain unexplained. LXRα and β are ligand-activated nuclear receptors that bind metabolites of cholesterol^18,19^. While LXRα/β is described in many reports as being an important pathway involved in many neurodegenerative diseases^36,37^, its specific role in CNS and PNS cells remains unclear and likely pleiotropic. Our current data suggest that LXRα/β may drive a unique transcriptional program regulating neuron-nerve-associated cells (likely SC) interactions that may sustain peripheral nerve function. Our findings using mice models (WD-fed and LXRα/β sensory neuron specific deletion) and *ex vivo* (DRG organotypic and primary neuron cultures), show that WD alters LXR activity that changes Nrg1 expression leading to neuron-SC communication impairment in the PNS. LXR activation had been shown to improve WD-induced neuropathy^16^. It is possible that LXR/Nrg1/ErbB pathway explain in part this improvement.

In the PNS, SC surrounds peripheral nerve axons with myelin membranes. Abnormal SC impairs PNS function^33,34,42^. During peripheral nerve development, SC differentiation and myelination critically depend on Nrg1, an axon-derived growth factor ^26-28^. Nrg1 belongs to a family of transmembrane and secreted epidermal growth factor (EGF)-like growth factors, which activates ErbB receptor tyrosine kinases^42^. When expressed on the neuron axonal surface, the transmembrane Nrg1-type III isoform regulates SC development and myelin sheath thickness^28,43,44^. Sensory neurons (both myelinated and non-myelinated) express different levels of Nrg1. The amount of Nrg1 present on the axonal membranes^23^ and SC myelin volume positively correlates with the surface area of the associated axon^24^. Data suggest that interaction between non-myelinating SC and small diameter axons in Remak bundles is coordinated by Nrg1^25,31^, indicating that both myelinating and non-myelinating SC and functionally modulated by Nrg1.

Nrg1 activation of ErbB receptors activate various signaling cascades which elicits cellular responses like proliferation, differentiation, motility, cell survival, and gene expression^23^. Both ErbB2 and ErbB3 receptors are required for signal transduction in the Schwann cells, but ErbB3, but not ErbB2, binds extracellular ligands with high affinity, however ErbB3 is catalytically inactive, and ErbB2 contributes to the tyrosine kinase activity essential for signaling^45^. Disruption of Nrg1/ErbB interaction leads to abnormal and aberrant myelination of large fibers and perturbation in the Remak bundle containing small non-myelinated axons resulting in abnormal nerve conduction and altered nerve structure^30,31^. Regulation of SC lipid synthesis via Srebf1 is one of the primary transcriptional responses following Nrg1-ErbB2/3 stimulation^46^. The mechanism of action of Nrg1 is not clear in non-myelinated fibers and needs to be better defined.

Evidence for the implication of LXR and oxysterols in the negative regulation of lipid and myelin gene expression was provided by Makoukji et al.,^36^. Using *in vitro* approaches, they showed that oxysterols are expressed in SC, and that they repress the gene expression of Mpz and Pmp22 by a mechanism involving LXR α/β^36^. The data are surprising because, although myelin gene transcripts are up-regulated in mice lacking LXR α/β, myelin sheath thickness is reduced *in vivo*^36^. They conclude that this hypomyelination in the LXR α/β knockout animals may be the consequence of an altered cholesterol homeostasis or an inefficient myelin protein trafficking from the endoplasmic reticulum^36^. Our current study using sensory neuron specific LXRα/β knockout establishes a role of LXR in sensory neurons-SC communication and provide another mechanism to explore and understand the role of LXR in the SC function.

Previous studies from others and our group showed that signaling pathway in the neurons of the DRG is disrupted inducing cellular stress^16,32,38,47^. Our previous data showed that activation of LXR might regulate lipid metabolism in sensory neurons of the DRG *in vivo* to blunt the WD-induced cellular stress in sensory neurons of the DRG^16^. Altogether, our data indicates that LXRs are active in the sensory neurons of the DRG where they regulate multiple pathways^16,36,37,40,41^ (cholesterol metabolism, ER stress pathway, membrane composition, neuron-SC communication via Nrg1). Further investigation would be necessary to better delineate how LXR regulates Nrg1 signal transduction. Sensory neuron and SC-specific studies would be helpful to decipher the role of lipids in peripheral neurons and better understand the complex Obesity-induced neuropathy. To this end, further investigations needs to better define the various SC in the PNS. After that, it would be important to evaluate the type of SC that interact with sensory neurons expressing LXR. For example, recently Abdo et al., unmasked a unique non-myelinating SC expressing Sox10 associated with small fiber neurons to regulate nociception^48^. It is possible that Nav1.8 and LXRs expressing axons are associated with Sox10-expressing SC to regulate PNS function. Cell specific studies and neuron-SC co-culture using appropriate markers - that still need to be characterized - will be necessary to specify new discoveries detailed in the current studies, however the markers and genetic models are currently lacking. This study will open-up avenues for future research aimed at understanding Obesity-associated neuropathies for which there is no cure yet; indeed restoring the communication between neurons and SC might be a new pathway to explore to treat and cure PNS dysfunction in Obesity-associated neuropathy.

## Materials and Methods

### Mice

All studies were conducted in accordance to recommendations in the Guide for the Care and Use of Laboratory Animals of the National Institutes of Health and the approval of the Loyola University Chicago Institutional Animal Care and Use Committee. C57BL/6J (#000664), RiboTag (#011029) were obtained from Jackson laboratory (Maine, USA) and crossed with transgenic mice carrying Cre recombinase driven by a Scn10a promoter (Nav1.8::Cre mice) to generate wild-type and RiboTag+/+:Nav1.8Cre+/- mice^16^. Sensory neuron specific liver x receptor (LXRα and β) knockouts were obtained by crossing LXRαfl/flβfl/fl mice with Nav1.8Cre+/- mice to generate LXRαfl/flβfl/fl:Nav1.8Cre+/- which were then crossed with LXRαfl/flβfl/fl to obtain LXRαfl/flβfl/fl (controls) and LXRαfl/flβfl/fl:Nav1.8Cre+/-^16,17^. All mice were housed 4/cage under a 12:12 h light/dark cycle. Mice received either NC (Teklad LM-485) or WD (TD88137, Teklad Diets; 42%kcal from fat, 34% sucrose by weight, and 0.2% cholesterol total) (Envigo, Indiana, USA) for 12 weeks starting at weaning^16,17^. All studies mentioned were done exclusively using male mice to avoid confounding effect of hormones with experimenter blinded to both treatment and genotype.

### In Situ Hybridization

Fluorescence in situ hybridization (FISH) was performed on 20μm thick slices of fresh frozen mouse DRG using RNAscope fluorescent multiplex reagents (Advanced Cell Diagnostics, 320850) according to the manufacturer’s instructions. RNA probes for Lxrα and Lxrβ (Advanced Cell Diagnostics, 440881 and 440871 respectively) were incubated with the DRG slices and signal amplification was achieved using the multiplex reagents as instructed. Images were captured using Olympus IX80 Inverted Microscope equipped with an X-Cite 120Q fluorescent light source (Lumen Dynamics) and a CoolSNAP HQ2 CD camera (Photometrics). Image processing was done using CellSens software (Olympus Corporation, Waltham, Massachusetts).

#### In vivo agonist treatment

WT and RiboTag mice were treated with vehicle or LXR agonist (GW3965; 25mg/kg BW) (Axon Medchem, Virginia, USA) by i.p. twice a week for 3 weeks starting at 8 weeks on WD as reported before^16^. Tissues were rapidly dissected and frozen in liquid nitrogen before analysis. Tissue from RiboTag mice were harvested and processed as detailed below.

#### DRG organotypic culture

Juvenile male mice (4-5weeks) were anesthetized with isoflurane before decapitation, and the DRG were quickly removed and cultured on a air-interface membrane (Millipore). Cultures were maintained for a week in standard culture medium^17^ replacing every other day in a 37°C and 5% CO_2_ incubator. After an overnight incubation in low serum (2.5%) MEM supplemented with GlutaMAX (2mM), DRG were stimulated with either vehicle or 15µM GW3965 for 24 hrs before palmitate treatment (400 µM) for another 24hrs as described previously^16^. RNA was extracted using Acturus PicoPure RNA Extraction Kit (Applied Biosystems, California, USA).

#### Enrichment of transcripts from sensory neurons

DRG from RiboTag+/+:Nav1.8Cre+/- mice were either freshly harvested for RNA isolation or harvested to perform organotypic culture followed by RNA isolation. To isolate RNA associated with HA-tagged ribosomes in sensory neurons, immunoprecipitation (IP) followed by mRNA purification following the procedure published by Sanz et al. was used^16,49^. Briefly, DRG were homogenized in homogenization buffer and supernatant removed after centrifuging at 10,000g for 10 min at 4°C. 10% of the homogenate was saved (input) for mRNA isolation. Remaining volume was incubated at 4°C with anti-HA antibody (Biolegend, #901513) at 1:150 dilution for 4hrs on a gentle spinner. This is followed by an overnight incubation at 4°C on a gentle spinner with above sample transferred to tube containing magnetic beads (Pierce A/G magnetic beads, California, USA). Supernatant form the samples were collected and beads were washed with high salt buffer, 3 times,10 min each at 4°C on spinner. After final wash, lysis buffer (RNeasy Micro Kit, Qiagen, Maryland, USA) with β-mercaptoethanol (10µl/ml) was added to elute the mRNA. Total RNA from the IP’ed polysomes was eluted using RNeasy Micro Kit (Qiagen, California, USA) following manufacturer’s instructions and quantified with Quant-iT RiboGreen RNA Assay kit (Invitrogen, California, USA) and Agilent Bioanalyzer. Quantitative PCR performed on cDNA reverse transcribed from Ribotag mice RNA were normalized to β-actin as previously reported^49-51^.

#### Primary DRG neuronal culture

DRG from juvenile male mice were collected in ice-cold advanced DMEM without any supplementation and axotomized as previously reported^16^. Briefly, axotomized DRG were then transferred to a collagenase A/trypsin mix (1.25mg/ml each) and incubated for 30min. Partially digested DRG were then passed through fire polished glass pipettes followed by 3min spin at 3000g. After careful removal of supernatant, cells were resuspended in advanced DMEM with 10% FBS and 4mM GlutaMAX, and plated onto a poly-l-lysine coated plates. Neuronal cultures were maintained in a 37°C and 5% CO_2_ incubator for 3-4 days changing above media supplemented with Ara-C (2µM) to inhibit replicative cells every other day before treating the cells to extract RNA as described above.

#### Luciferase Reporter Assay

Primary DRG neuronal cultures isolated as described above were transduced with pGreenFire1-LXRE in LXRα Lentivector (System Biosciences, California) followed by respective group treatment 48hrs after transduction. pGreenFire1-LXRE in LXRα co-expresses a destabilized copepod GFP and luciferase from the LXR response elements and neighboring regions in the LXRα promoter paired with a mCMV promoter. After 24hours of treatment, cells were lysed in equal volume of ONE-Glo Lysis Buffer (Promega, Wisconsin) to that of culture medium. The luminescence was measured using CLARIOstar (BMG Labtech) and normalized to total protein.

### RNA isolation, cDNA library construction and Illumina sequencing

Total RNA was extracted from Sciatic nerve of NC- or WD-fed mice with Arcturus PicoPure RNA isolation kit (Applied Biosystems). Two biological replicates were used for each group. Total RNA was quantified by Qubit and assessed for quality on an Agilent Bioanalyzer using Total RNA Pico Chip. Total RNA samples that passed QC were used as input for library construction. Full-length cDNA synthesis and amplification were carried out with the Clontech SMART-Seq v4 Ultra Low Input RNA Kit. Subsequently, Illumina sequencing libraries were prepared from the amplified full-length cDNA with the Nextera XT DNA Library Preparation Kit. Prior to sequencing, the prepared libraries were quantified with Qubit and validated on a Bioanalyzer with a High Sensitivity DNA chip. The sequencing of the libraries was conducted on an Illumina NextSeq 500 NGS System. Single 75 bp reads were generated with dual indexing. RNA-seq Analysis done with STAR and DESeq2. The quality of reads, in FASTQ format, was evaluated using FastQC. Reads were trimmed to remove Illumina adapters from the 3’ ends using cutadapt^52^. Trimmed reads were aligned to the Mus musculus genome (mm10) using STAR^53^. Read counts for each gene were calculated using htseq-count^54^ in conjunction with a gene annotation file for mm10 obtained from Ensembl (http://useast.ensembl.org/index.html). Normalization and differential expression were calculated using DESeq2 that employs the Wald test^55^. The cutoff for determining significantly differentially expressed genes was an FDR-adjusted p-value less than 0.05 using the Benjamini-Hochberg method.

#### Validation of transcripts and Quantitative PCR

For all genes of interest, qPCR was performed using Sybr green-based assay (Roche, Indiana, USA) using IDT primers (IDT technologies, Iowa, USA). 18s (β-actin, for RiboTag:Nav1.8 IP’ed mRNA) was used to normalize data and quantification was done using ΔΔCT method with vehicle treated group’s mean value set at 100% as reported before^16^.

#### Quantification and statistical analysis

All data are represented as mean±S.E.M. Analysis were done using IBM SPSS Statistics 24. For single group comparisons either a 1- or 2-tailed t-test was used as appropriate and multiple comparisons were performed using ANOVA. For repeated measures, 2-way ANOVA was used and p value less than 0.05 was considered significant. Number of experiments/replicates and mice for each experiment are described in figure legends.

## Author contributions

CKG and VM-A were involved in the conception and design of the experiments. CKG, VM-A, and RB were involved in data collection, assembly, analysis and interpretation of data. CKG and VM-A drafted the manuscript. CKG, VM-A, and RB reviewed the manuscript.

## Acknowledgements and funding

We thank Loyola University Chicago animal facility for mice housing. We acknowledge the service of NUSeq Core (Northwestern University, Chicago). This work was supported by Loyola University Chicago and NIH R01 DK117404 to VM-A.

## References

1 Martini, R., Fischer, S., López-Vales, R. & David, S. Interactions between Schwann cells and macrophages in injury and inherited demyelinating disease. Glia 56, 1566–1577, doi:10.1002/glia.20766 (2008).

2 Martini, R. & Willison, H. Neuroinflammation in the peripheral nerve: Cause, modulator, or bystander in peripheral neuropathies? Glia 64, 475–486, doi:10.1002/glia.22899 (2016).

3 Feldman, E. L. et al. Diabetic neuropathy. Nature Reviews Disease Primers 5, 41, doi:10.1038/s41572-019-0092-1 (2019).

4 Pop-Busui, R. et al. DCCT and EDIC studies in type 1 diabetes: lessons for diabetic neuropathy regarding metabolic memory and natural history. Curr Diab Rep 10, 276–282, doi:10.1007/s11892-010-0120-8 (2010).

5 Ang, L., Jaiswal, M., Martin, C. & Pop-Busui, R. Glucose control and diabetic neuropathy: lessons from recent large clinical trials. Curr Diab Rep 14, 528, doi:10.1007/s11892-014-0528-7 (2014).

6 Pop-Busui, R. et al. Impact of glycemic control strategies on the progression of diabetic peripheral neuropathy in the Bypass Angioplasty Revascularization Investigation 2 Diabetes (BARI 2D) Cohort. Diabetes Care 36, 3208–3215, doi:10.2337/dc13-0012 (2013).

7 Wiggin, T. D. et al. Elevated triglycerides correlate with progression of diabetic neuropathy. Diabetes 58, 1634–1640, doi:10.2337/db08-1771 (2009).

8 Hur, J. et al. Transcriptional networks of murine diabetic peripheral neuropathy and nephropathy: common and distinct gene expression patterns. Diabetologia 59, 1297-1306, doi:10.1007/s00125-016-3913-8 (2016).

9 Pande, M. et al. Transcriptional profiling of diabetic neuropathy in the BKS db/db mouse: a model of type 2 diabetes. Diabetes 60, 1981–1989, doi:10.2337/db10-1541 (2011).

10 Tashani, O. A., Astita, R., Sharp, D. & Johnson, M. I. Body mass index and distribution of body fat can influence sensory detection and pain sensitivity. Eur J Pain, doi:10.1002/ejp.1019 (2017).

11 Jin, H. Y. & Park, T. S. Role of inflammatory biomarkers in diabetic peripheral neuropathy. J Diabetes Investig 9, 1016–1018, doi:10.1111/jdi.12794 (2018).

12 Pop-Busui, R., Ang, L., Holmes, C., Gallagher, K. & Feldman, E. L. Inflammation as a Therapeutic Target for Diabetic Neuropathies. Curr Diab Rep 16, 29, doi:10.1007/s11892-016-0727-5 (2016).

13 Vincent, A. M., Calabek, B., Roberts, L. & Feldman, E. L. Biology of diabetic neuropathy. Handb Clin Neurol 115, 591–606, doi:10.1016/B978-0-444-52902-2.00034-5 (2013).

14 Arvidson, B. A study of the perineurial diffusion barrier of a peripheral ganglion. Acta Neuropathol 46, 139–144, doi:10.1007/bf00684815 (1979).

15 Liu, C. et al. PPARγ in vagal neurons regulates high-fat diet induced thermogenesis. Cell Metab 19, 722–730, doi:10.1016/j.cmet.2014.01.021 (2014).

16 Gavini, C. K. et al. Liver X Receptors Protect Dorsal Root Ganglia from Obesity-Induced Endoplasmic Reticulum Stress and Mechanical Allodynia. Cell Rep 25, 271-277.e274, doi:10.1016/j.celrep.2018.09.046 (2018).

17 Mansuy-Aubert, V. et al. Loss of the liver X receptor LXRα/β in peripheral sensory neurons modifies energy expenditure. Elife 4, doi:10.7554/eLife.06667 (2015).

18 Chawla, A., Repa, J. J., Evans, R. M. & Mangelsdorf, D. J. Nuclear receptors and lipid physiology: opening the X-files. Science 294, 1866–1870, doi:10.1126/science.294.5548.1866 (2001).

19 Repa, J. J. & Mangelsdorf, D. J. Nuclear receptor regulation of cholesterol and bile acid metabolism. Curr Opin Biotechnol 10, 557–563, doi:10.1016/s0958-1669(99)00031-2 (1999).

20 Livesey, F. J. et al. A Schwann cell mitogen accompanying regeneration of motor neurons. Nature 390, 614–618, doi:10.1038/37615 (1997).

21 Guénard, V., Gwynn, L. A. & Wood, P. M. Astrocytes inhibit Schwann cell proliferation and myelination of dorsal root ganglion neurons in vitro. J Neurosci 14, 2980–2992 (1994).

22 Stallcup, W. B., Arner, L. S. & Levine, J. M. An antiserum against the PC12 cell line defines cell surface antigens specific for neurons and Schwann cells. J Neurosci 3, 53–68 (1983).

23 Newbern, J. & Birchmeier, C. Nrg1/ErbB signaling networks in Schwann cell development and myelination. Seminars in cell & developmental biology 21, 922–928, doi:10.1016/j.semcdb.2010.08.008 (2010).

24 Smith, K. J., Blakemore, W. F., Murray, J. A. & Patterson, R. C. Internodal myelin volume and axon surface area. A relationship determining myelin thickness? J Neurol Sci 55, 231–246, doi:10.1016/0022-510x(82)90103-4 (1982).

25 Chen, S. et al. Disruption of ErbB receptor signaling in adult non-myelinating Schwann cells causes progressive sensory loss. Nature Neuroscience 6, 1186–1193, doi:10.1038/nn1139 (2003).

26 Taveggia, C. et al. Neuregulin-1 type III determines the ensheathment fate of axons. Neuron 47, 681–694, doi:10.1016/j.neuron.2005.08.017 (2005).

27 Syed, N. et al. Soluble neuregulin-1 has bifunctional, concentration-dependent effects on Schwann cell myelination. J Neurosci 30, 6122–6131, doi:10.1523/JNEUROSCI.1681-09.2010 (2010).

28 Nave, K. A. & Salzer, J. L. Axonal regulation of myelination by neuregulin 1. Curr Opin Neurobiol 16, 492–500, doi:10.1016/j.conb.2006.08.008 (2006).

29 Zanazzi, G. et al. Glial growth factor/neuregulin inhibits Schwann cell myelination and induces demyelination. J Cell Biol 152, 1289–1299, doi:10.1083/jcb.152.6.1289 (2001).

30 Chen, S. et al. Neuregulin 1-erbB signaling is necessary for normal myelination and sensory function. J Neurosci 26, 3079–3086, doi:10.1523/JNEUROSCI.3785-05.2006 (2006).

31 Fricker, F. R. et al. Sensory axon-derived neuregulin-1 is required for axoglial signaling and normal sensory function but not for long-term axon maintenance. J Neurosci 29, 7667–7678, doi:10.1523/JNEUROSCI.6053-08.2009 (2009).

32 Jayaraj, N. D. et al. Reducing CXCR4-mediated nociceptor hyperexcitability reverses painful diabetic neuropathy. J Clin Invest 128, 2205–2225, doi:10.1172/JCI92117 (2018).

33 Jessen, K. R. & Mirsky, R. The repair Schwann cell and its function in regenerating nerves. The Journal of physiology 594, 3521–3531, doi:10.1113/JP270874 (2016).

34 Brown, A. M., Evans, R. D., Black, J. & Ransom, B. R. Schwann cell glycogen selectively supports myelinated axon function. Annals of neurology 72, 406–418, doi:10.1002/ana.23607 (2012).

35 Cermenati, G. et al. Lack of sterol regulatory element binding factor-1c imposes glial Fatty Acid utilization leading to peripheral neuropathy. Cell Metab 21, 571–583, doi:10.1016/j.cmet.2015.02.016 (2015).

36 Makoukji, J. et al. Interplay between LXR and Wnt/β-catenin signaling in the negative regulation of peripheral myelin genes by oxysterols. J Neurosci 31, 9620–9629, doi:10.1523/JNEUROSCI.0761-11.2011 (2011).

37 Wang, L. et al. Liver X receptors in the central nervous system: from lipid homeostasis to neuronal degeneration. Proc Natl Acad Sci U S A 99, 13878–13883, doi:10.1073/pnas.172510899 (2002).

38 Vincent, A. M. et al. Dyslipidemia-induced neuropathy in mice: the role of oxLDL/LOX-1. Diabetes 58, 2376–2385, doi:10.2337/db09-0047 (2009).

39 Haskins, J. W. et al. Neuregulin-activated ERBB4 induces the SREBP-2 cholesterol biosynthetic pathway and increases low-density lipoprotein uptake. Sci Signal 8, ra111, doi:10.1126/scisignal.aac5124 (2015).

40 Hong, C. & Tontonoz, P. Coordination of inflammation and metabolism by PPAR and LXR nuclear receptors. Curr Opin Genet Dev 18, 461–467, doi:10.1016/j.gde.2008.07.016 (2008).

41 Rong, X. et al. LXRs regulate ER stress and inflammation through dynamic modulation of membrane phospholipid composition. Cell Metab 18, 685–697, doi:10.1016/j.cmet.2013.10.002 (2013).

42 Fledrich, R. et al. NRG1 type I dependent autoparacrine stimulation of Schwann cells in onion bulbs of peripheral neuropathies. Nature Communications 10, 1467, doi:10.1038/s41467-019-09385-6 (2019).

43 Birchmeier, C. & Nave, K. A. Neuregulin-1, a key axonal signal that drives Schwann cell growth and differentiation. Glia 56, 1491–1497, doi:10.1002/glia.20753 (2008).

44 Michailov, G. V. et al. Axonal neuregulin-1 regulates myelin sheath thickness. Science 304, 700–703, doi:10.1126/science.1095862 (2004).

45 Citri, A., Skaria, K. B. & Yarden, Y. The deaf and the dumb: the biology of ErbB-2 and ErbB-3. Exp Cell Res 284, 54–65, doi:10.1016/s0014-4827(02)00101-5 (2003).

46 Nave, K. A. Myelination and support of axonal integrity by glia. Nature 468, 244–252, doi:10.1038/nature09614 (2010).

47 Menichella, D. M. et al. CXCR4 chemokine receptor signaling mediates pain in diabetic neuropathy. Mol Pain 10, 42, doi:10.1186/1744-8069-10-42 (2014).

48 Abdo, H. et al. Specialized cutaneous Schwann cells initiate pain sensation. Science 365, 695–699, doi:10.1126/science.aax6452 (2019).

49 Sanz, E. et al. Cell-type-specific isolation of ribosome-associated mRNA from complex tissues. Proc Natl Acad Sci U S A 106, 13939–13944, doi:10.1073/pnas.0907143106 (2009).

50 Nectow, A. R. et al. Rapid Molecular Profiling of Defined Cell Types Using Viral TRAP. Cell Rep 19, 655–667, doi:10.1016/j.celrep.2017.03.048 (2017).

51 Soden, M. E. et al. Genetic Isolation of Hypothalamic Neurons that Regulate Context-Specific Male Social Behavior. Cell Rep 16, 304–313, doi:10.1016/j.celrep.2016.05.067 (2016).

52 Martin, M. Cutadapt removes adapter sequences from high-throughput sequencing reads. EMBnet.journal; Vol 17, No 1: Next Generation Sequencing Data AnalysisDO - 10.14806/ej.17.1.200 (2011).

53 Dobin, A. et al. STAR: ultrafast universal RNA-seq aligner. Bioinformatics (Oxford, England) 29, 15–21, doi:10.1093/bioinformatics/bts635 (2013).

54 Anders, S., Pyl, P. T. & Huber, W. HTSeq--a Python framework to work with high-throughput sequencing data. Bioinformatics (Oxford, England) 31, 166–169, doi:10.1093/bioinformatics/btu638 (2015).

55 Love, M. I., Huber, W. & Anders, S. Moderated estimation of fold change and dispersion for RNA-seq data with DESeq2. Genome biology 15, 550–550, doi:10.1186/s13059-014-0550-8 (2014).

